# *Staphylococcus aureus* lipid factors modulate melanoma clustering and invasion in zebrafish

**DOI:** 10.1101/2024.02.27.582365

**Authors:** Morgan A. Giese, Gayathri Ramakrishnan, Laura H. Steenberge, Jerome X. Dovan, John-Demian Sauer, Anna Huttenlocher

**Affiliations:** Department of Medical Microbiology and Immunology, University of Wisconsin-Madison, Madison, WI, USA; Cellular and Molecular Biology Graduate Program, University of Wisconsin-Madison, Madison, WI, USA; Cancer Biology Graduate Program, University of Wisconsin-Madison, Madison, WI, USA; Department of Biochemistry, University of Wisconsin-Madison, Madison, WI, USA; University of Wisconsin School of Medicine and Public Health, Madison, Wisconsin, USA; Morgridge Institute for Research, Madison, Wisconsin, USA; University of Wisconsin Medical Scientist Training Program (MSTP) Summer Scholars, University of Wisconsin-Madison, Madison, WI, USA; Department of Nutritional Sciences, University of Wisconsin-Madison, Madison, WI, USA.; Department of Pediatrics, University of Wisconsin-Madison, Madison, WI, USA

**Keywords:** melanoma, zebrafish, *S. aureus*, lipids, skin microbiome

## Abstract

The microbiome can influence cancer development and progression. However, less is known about the role of the skin microbiota in melanoma. Here, we take advantage of a zebrafish melanoma model to probe the effects of *Staphylococcus aureus* on melanoma invasion. We find that *S. aureus* produces factors that enhance melanoma invasion and dissemination in zebrafish larvae. We used a published *in vitro* 3D cluster formation assay which correlates increased clustering with tumor invasion. *S. aureus* supernatant increased clustering of melanoma cells which was abrogated by use of a Rho-Kinase inhibitor, implicating a role for Rho-GTPases. The melanoma clustering response was specific to *S. aureus* and not other staphylococcal species, including *S. epidermidis*. Our findings suggest that *S. aureus* may promote melanoma clustering and invasion via lipids generated by the lipase SAL2 (*gehB*). Taken together, these findings suggest that specific bacterial products mediate melanoma invasive migration in zebrafish.

**Summary statement:** *S. aureus* supernatant increases melanoma invasion in zebrafish and is likely mediated by production of bacterial lipids.

## Introduction

The host microbiome is capable of influencing all aspects of health including cancer initiation and progression [1, 2]. Oncomicrobes, or microbes with carcinogenic properties, cause an estimated 2.2 million cancer cases per year [3]. Furthermore, there is a growing list of “complicit” microbes which are capable of promoting cancer progression [4]. These microbes can alter proliferative versus cell death signals, produce DNA damaging toxins, or induce cancer cell invasion by triggering epithelial-to-mesenchymal transition (EMT) [5-7]. Additionally, microbes are highly capable of modulating the immune response, either promoting an inflammatory tumor microenvironment or an immunosuppressive environment that prevents tumor cell killing [4, 5, 8]. As the gut is the largest reservoir of microbes in the human body, most studies have focused on the impact of gut microbes on cancer development and therapy response [9, 10]. The skin represents the second largest microbiota population in the body, yet few studies have evaluated the role that these microbes play in cancer development.

Increasing evidence indicates that skin microbes, such as *Staphylococcus aureus,* are linked to cancers including cutaneous T cell lymphoma [11-13]. *S. aureus* is commonly found on healthy skin and colonizes 20-40% of the general population [14], yet its presence is a leading risk for surgical site infections as *S. aureus* can transition to a pathogenic state with changes in the environment [15]. Non-melanoma tumors were found to have an overabundance of *S. aureus* compared to healthy skin [16, 17], and challenge with *S. aureus* increased human Squamous Cell Carcinoma (SCC) proliferation [18]. For melanoma, only one publication has profiled the microbial community on patients and found that *Propionibacterium*, *Staphylococcus* and *Corynebacterium* were most the common genera on both healthy skin and melanocytic lesions [19]. In pig skin models, *Staphylococcus* species were more prevalent on cutaneous melanoma compared to healthy skin [20], but the effect of these bacteria on the melanoma microenvironment is unknown. Melanoma occurrence has steadily increased in the United States, with 100,000 predicted diagnoses in 2023, and accounts for the majority of skin cancer related deaths (*American Cancer Society*). Given the correlation between *S. aureus* and the development of other skin cancers, we sought to determine the effect of *S. aureus* on cutaneous melanoma growth and progression.

Here we used a larval zebrafish model to investigate the impact of *S. aureus* on melanoma progression. Zebrafish models for cancer cell transplantation are well developed and allow for study of the early stages of cancer invasion [21-24]. In this study, we show that *S. aureus* produces factors capable of enhancing melanoma cell invasion and dissemination. To determine the mechanism driving melanoma invasion, we took advantage of an *in vitro* 3D cluster formation assay using low attachment plates. More invasive melanoma cells were previously shown to form large spherical clusters due to increased expression of adhesion genes [25]. We found that incubation with *S. aureus* supernatant increased clustering of melanoma cells, which was abrogated by use of the Rho-Kinase inhibitor (ROCK). The clustering response was specific to *S. aureus* and not other staphylococcal species, including *S. epidermidis*, or other gram-negative bacteria tested. Furthermore, we determined that *S. aureus* may promote cluster formation via lipids generated by the lipase SAL2 (*gehB*), as genetic mutation resulted in reduced melanoma clustering and invasion. Therefore, these findings suggest that *S. aureus* produces lipids that promote melanoma invasion.

## Results

### *S. aureus* supernatant promotes melanoma invasion and dissemination *in vivo*

To evaluate the effect of *S. aureus* on melanoma cell behavior *in vivo*, ZMEL1-GFP zebrafish melanoma cells were incubated with *S. aureus* bacterial supernatant for three days and then injected into the larval zebrafish hindbrain (Figure 1A). Bacterial supernatants were used over live bacteria to limit bacterial overgrowth. The larval hindbrain has previously been used as a model to assess melanoma and other cancer cell dissemination *in vivo* [21, 24]. Larval zebrafish are optically transparent which allows for *in vivo* visualization of cell behavior, including local cancer cell invasion and dissemination [26, 27]. Using fluorescent confocal imaging, we visualized melanoma invasion over time. Two days post injection (2dpi), melanoma invasion away from the cell mass was increased after pre-incubation with bacterial supernatant, compared to culture in media alone (Figure 1B,C). Furthermore, ZMEL1 cells displayed an elongated morphology suggestive of the mesenchymal cell migration of invading tumor cells (Figure 1B) [28, 29]. Following migration into the tissue, cancer cells can invade into the skin or enter the vasculature to spread to other regions of the zebrafish [25]. We imaged cancer cell dissemination in the tail fin and found that melanoma cells cultured in *S. aureus* supernatant were significantly more likely to disseminate (Figure 1D). Our findings show that pre-treatment of melanoma cells with *S. aureus* supernatant modifies melanoma cell behavior and increases invasion and dissemination into tissues.

**Figure 1.**
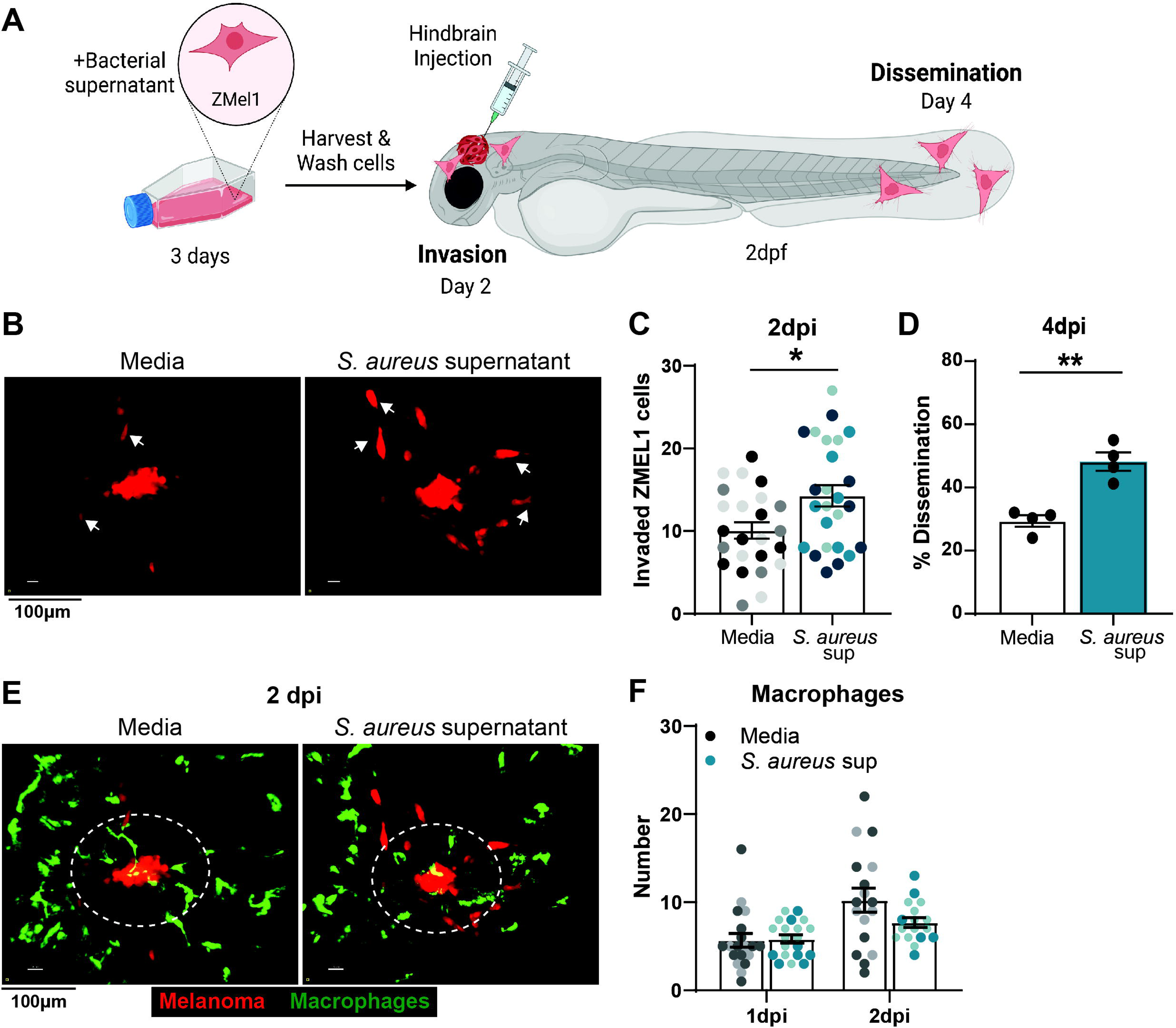
*S. aureus* supernatant promotes melanoma invasion and dissemination *in vivo*. (A) Schematic of ZMEL1-td-Tomato injection into larval zebrafish hindbrain. Confocal imaging was performed in the hindbrain at 2 days post injection (dpi) and in the tail at 4dpi. (B) Representative images and (C) quantification of ZMEL1 invasion at 2dpi in the zebrafish hindbrain after culture in media alone or *S. aureus* USA300 supernatant. From 3 independent experiments. n=24-25 larvae per condition. Arrows indicate migrated melanoma cells with an elongated morphology. (D) Percent of zebrafish with disseminated ZMEL1 cells in the tailfin at 4dpi. n=130 larvae for media control, n=119 larvae with *S. aureus* supernatant. (E) Representative images at 2dpi and (F) quantification of macrophage recruitment to ZMEL1 melanoma cells. n=17-20 larvae per condition. Data represents two replicates. Diagram (A) was created with BioRender.com. Experiments (C, D) were conducted three times and dots represent independent larvae color coded per replicate. Dots in (D) represent independent replicates. Bars indicate the mean ± SEM. Dotted circles represent 50um region where recruited immune cells were counted. *p* values were calculated by unpaired T-test (C) or paired T-test (D). *, p < 0.05; **, p < 0.01

We next wanted to determine if increased melanoma cell invasion was associated with increased immune infiltration. Increased recruitment of immune cells can promote cancer cell progression and metastasis [30, 31]. Larval zebrafish have an intact innate immune system which can be live imaged to evaluate immune cell infiltration and association with tumor cells [27, 32]. Using fluorescent reporters for neutrophils and macrophages, we imaged innate immune cell recruitment to the injection site. Neutrophils were recruited early, arriving within hours of melanoma injection, but showed little interest in ZMEL1 cells. We found no difference in the number of neutrophils with *S. aureus* supernatant incubation versus media alone (Figure S1A, B). Macrophages are secondarily recruited and in greater numbers with peak recruitment at 2 dpi. Similar to neutrophils, we found no significant difference in the level of macrophage recruitment with melanoma cells pretreated with *S. aureus* supernatant (Figure 1E,F). Therefore, ZMEL1 cell invasion and dissemination to the fin is likely due to direct effects of the supernatants on melanoma cells.

### *S. aureus* supernatant promotes melanoma clustering *in vitro*

To determine how *S. aureus* supernatant promotes melanoma invasion, we used a previously developed 3D *in vitro* clustering assay on low attachment plates [25]. Increased invasion and metastasis of ZMEL1 melanoma cells in zebrafish was shown to correlate with enhanced *in vitro* clustering response due to increased cell-cell adhesion [25]. We tested the effect of *S. aureus* supernatant on melanoma cell clustering by imaging cells over the course of 7 days. We found little difference in cluster size for the first 3 days of culture. However, after day 4, melanoma cell clusters exponentially increased in size when cultured in the presence of *S. aureus* bacterial supernatant (Figure 2A, B). At 7 days, *S. aureus* supernatant significantly increased melanoma cluster size compared to media alone (Figure 2C). To evaluate if *S. aureus* bacteria could promote clustering, we co-cultured melanoma with equivalent CFUs of heat killed *S. aureus*. We found a small increase in clustering over media alone, but this was significantly reduced compared to supernatant (Figure 2C). In summary, *S. aureus* supernatant is capable of producing secreted factors which promote melanoma clusters *in vitro*, correlating with the increased invasion observed *in vivo*.

**Figure 2.**
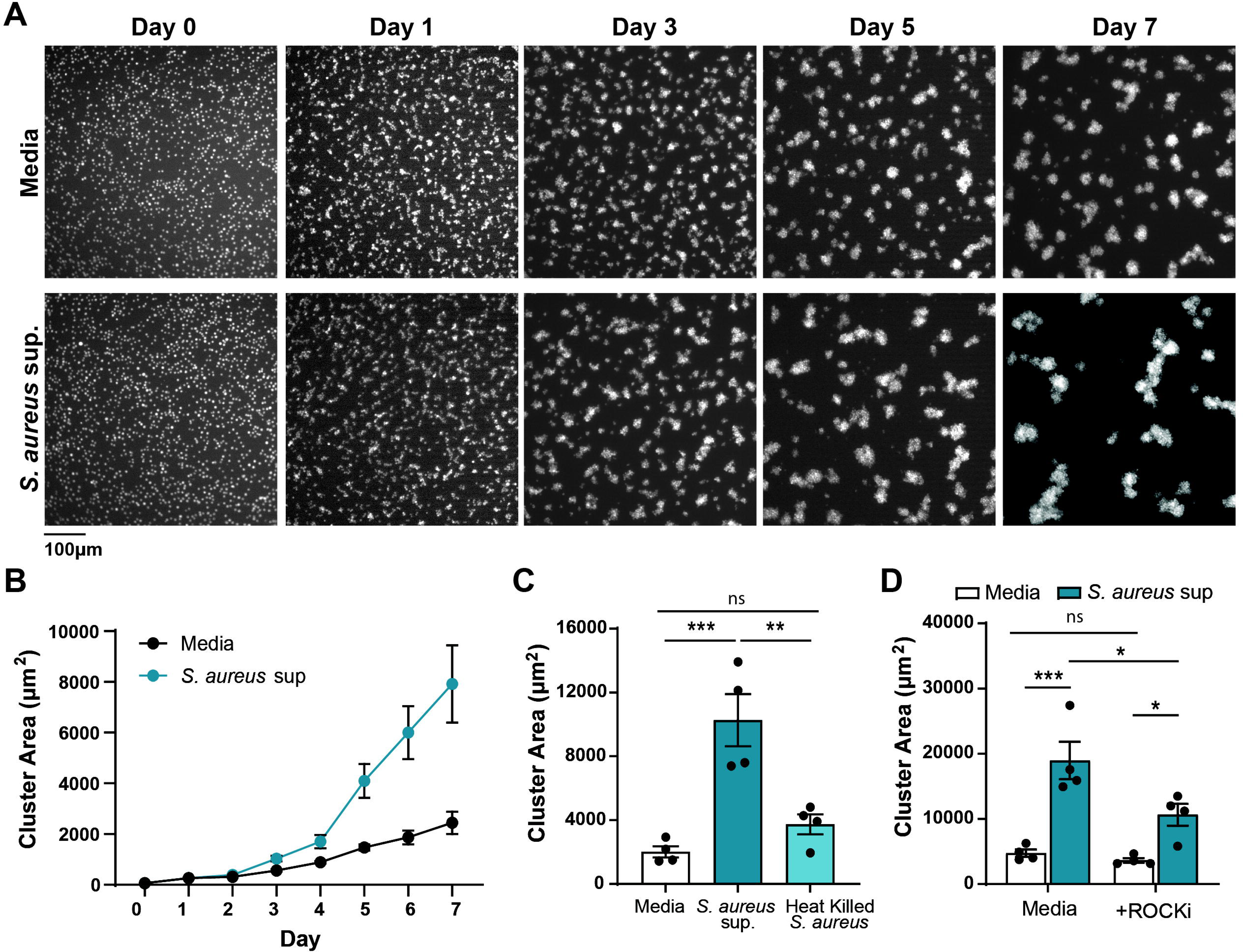
*S. aureus* supernatant promotes melanoma clustering *in vitro*. (A) Representative images of ZMEL1-EGFP melanoma cells cultures in the presence of media alone or *S. aureus* USA300 supernatant imaged over 7 days. (B) Plot shows one representative experiment with quantification of melanoma cluster area (µm^2^) over time. (C) Melanoma cluster size (µm^2^) with *S. aureus* USA300 supernatant or heat killed bacteria at 7 days. (D) Melanoma cluster size (µm^2^) with *S. aureus* USA300 supernatant plus 500nM ROCK inhibitor Y-27632 at 7 days. Experiments were conducted at least three times. Dots in (B) represent mean ± SEM. Dots in (C, D) represent independent replicates and bars indicate the mean ± SEM. *p* values were calculated by One-way ANOVA, (C) or Two-way ANOVA (D). *, p < 0.05; **, p < 0.01; ***, p < 0.001.

Next, we live imaged melanoma cells over time to determine if increased cluster size was due to enhanced cell-cell adhesion, as previously described [25]. Melanoma cells cultured in bacterial supernatant were more active than cells in media alone and clusters appeared to migrate towards one another to promote larger cluster formation (Movie 1, Movie 2). To determine if *S. aureus* supernatant promotes cluster formation due to increased melanoma cell migration, we utilized a Rho-Kinase (ROCK) inhibitor Y27632. ROCK acts downstream of Rho GTPase to promote cytoskeletal rearrangement and cell motility [33]. Furthermore, Rho signaling can promote cancer cell migration and invasion [34]. Addition of ROCK inhibitor to the melanoma cells cultured with bacterial supernatant significantly diminished cluster formation (Figure 2D). There was no effect of ROCK inhibitor on melanoma cells incubated with media control, indicating that the ability to cluster was not affected. Therefore, our findings suggest that *S. aureus* supernatant promotes melanoma cell migration via activation of Rho-GTPase signaling.

### Melanoma clustering is specific to *S. aureus* species

We next determined if melanoma clustering is specific to *S. aureus* USA300 strain or can be induced by a broader spectrum of bacteria. Gram positive bacteria, such as *S. aureus*, are recognized by melanoma cells through the toll-like receptor TLR2 [35, 36]. We added the TLR2 agonist Pam3CSK4, but found no effect on melanoma cluster size indicating that TLR2 activation alone is not sufficient to promote cluster formation (Figure 3A). Next, we tested a second strain *S. aureus* Newman, which is a methicillin sensitive *Staphylococcus aureus* strain (MSSA) with high genetic similarity to USA300 [37]. Newman was capable of promoting melanoma cluster formation with levels similar to USA300 (Figure 3A). On the contrary, the commensal species *S. epidermidis* did not induce clusters (Figure 3A), suggesting that not all *Staphylococcus* species are capable of promoting melanoma clustering.

**Figure 3.**
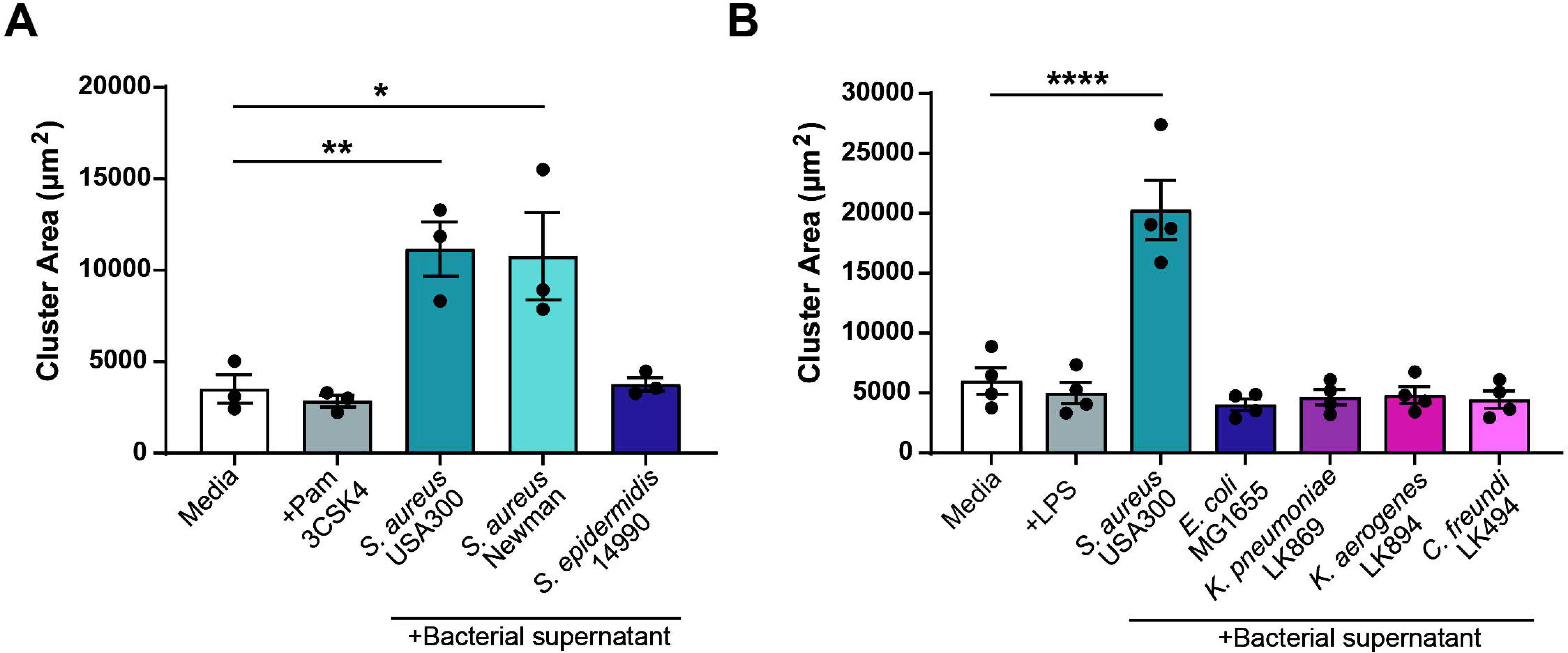
Melanoma clustering is specific to *S. aureus* species. Quantification of ZMEL1 melanoma cell cluster size (µm^2^) after 7 days of culture in media alone or plus bacterial supernatant. (A) Pam3CSK4 (100ug/mL) or bacterial supernatants from a selection of gram-positive bacteria were added to melanoma cell culture. (B) LPS (1ug/mL) or bacterial supernatants from a selection of gram-negative bacteria were added to melanoma cell culture. Experiments were conducted at least three times. Dots represent independent replicates and bars indicate the mean ± SEM. *p* values were calculated by One-way ANOVA. *, p < 0.05; **, p < 0.01; ****, p < 0.0001.

We also tested a selection of gram negative species including 3 clinical isolates from human skin (*K. pneumoniae, K. aerogenes, C. freundi*). None of the tested gram negative bacteria were capable of inducing melanoma cluster formation (Figure 3B). Furthermore, addition of the TLR4 agonist LPS did not affect melanoma clustering. Taken together, these findings indicate that the effect of bacterial supernatant on melanoma clustering is specific to the *S. aureus* species tested and not induced by TLR activation alone.

### *S. aureus* effect on melanoma clustering is likely mediated by lipids

Next, we modified the culture media used to grow *S. aureus* to determine the effect of supplemented media components on melanoma clustering. Microbiota utilize available host components during infection, including *S. aureus* which incorporates human serum lipids into its membrane during pathogenesis [38]. Thus, addition of host nutrients may influence production of the clustering factor by *S. aureus*. We generated supernatants from *S. aureus* grown in RPMI alone or supplemented with fetal bovine serum (FBS) or bovine serum albumin (BSA). While there was a trend towards larger clusters with the addition of FBS over serum free media, this was not significant (Figure 4A). These data suggest that serum components do not affect *S. aureus* production of the clustering factor. Interestingly, addition of BSA to bacterial supernatant significantly enlarged melanoma cluster size, with almost a 3-fold increase (Figure 4A). BSA is often included as a media supplement in serum free media as it promotes cell growth and survival. It does so by binding essential media components including fatty acids and other lipids to increase their concentration and interaction and uptake into cells [39, 40]. The increased clustering with BSA indicates that albumin may be able to bind hydrophobic molecules produced by *S. aureus* to enhance melanoma clustering. Furthermore, albumin proteins are present in FBS and may account for the slight increase in cluster size we observed with FBS supplementation.

**Figure 4.**
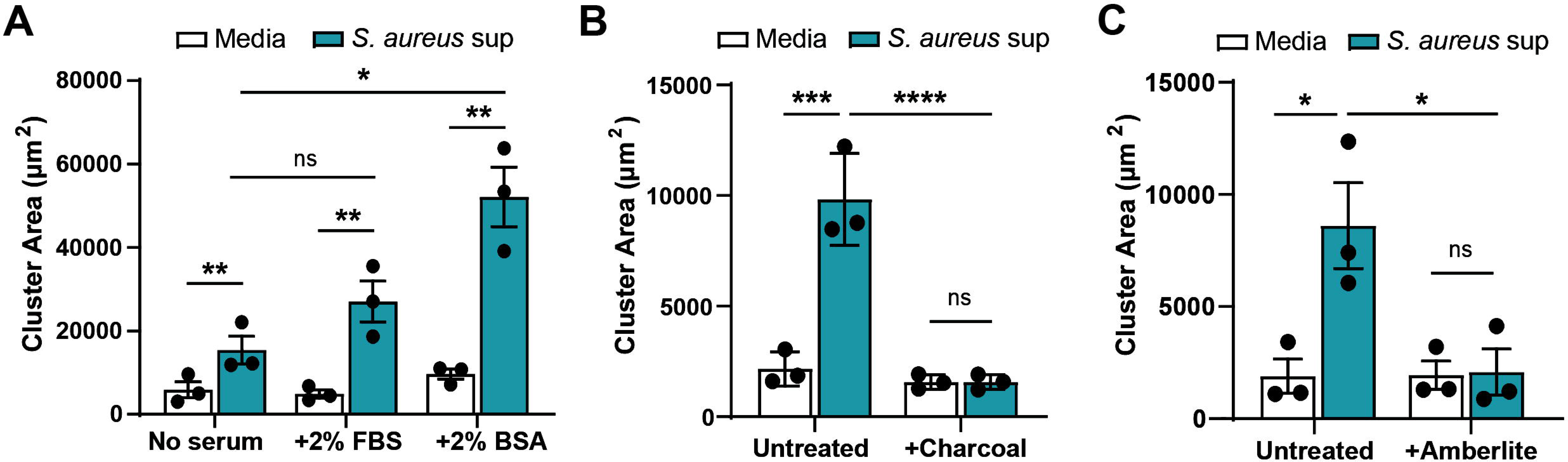
*S. aureus* effect on melanoma clustering is likely mediated by lipids. Quantification of ZMEL1 melanoma cells cluster size (µm^2^) after 7 days of culture in media alone or plus *S. aureus* USA300 bacterial supernatant. (A) Supernatants were collected from *S. aureus* USA300 bacteria grown in RPMI media alone or media supplemented with 2%FBS or 2%BSA, then cultured with melanoma cells. Supernatants collected from *S. aureus* USA300 bacteria grown in RPMI media alone were treated with (B) charcoal-dextran or (C) Amberlite-XAD4 and cultured with melanoma cells. Experiments were conducted at least three times. Dots represent independent replicates and bars indicate the mean ± SEM. *p* values were calculated by Two-way ANOVA. **, p < 0.01; ***, p < 0.001; ****, p < 0.0001.

To test if hydrophobic molecules in *S. aureus* supernatant are responsible for melanoma clustering, we stripped the bacterial supernatant. Stripping with charcoal-dextran completely abrogated melanoma clustering (Figure 4B). Use of a second stripping method with Amberlite XAD4 beads, which are highly adsorbent for hydrophobic compounds, also resulted in loss of the clustering phenotype (Figure 4C). We confirmed charcoal stripping of *S. aureus* lipids using TLC and found that polar, but not non-polar lipids were selectively depleted (Figure S2A, B). Thus, a hydrophobic molecule in *S. aureus* supernatant likely mediates clustering. Our data suggest these factors may be polar lipids.

### Deletion of *S. aureus* lipases alters melanoma clustering

Bacteria synthesize lipids necessary for key cell functions including formation of the cellular envelope [38, 41]. Production of these lipids is mediated by a cascade of enzymes [42]. Taking advantage of an available *S. aureus* USA300 transposon mutant library, we tested the ability of lipase mutants to promote melanoma clustering. *S. aureus* secrete a variety of lipases including SAL2, also known as GehB (SAUSA300_0320). Sal2 has ester hydrolase activity and cleaves both short and long chain triglycerides [43]. We found that supernatant from *S. aureus* ΔSAL2 significantly decreased cluster size (Figure 5A), indicating that SAL2 activity produces lipids that promote clustering. Sequence alignment to *S. aureus* found no SAL2 ortholog in *E. coli*, *K. pneumoniae,* or *K. aerogenes* [44], correlating to our clustering data. While *S. epidermidis* contains a SAL2 ortholog [45], its level of secretion is unclear.

**Figure 5.**
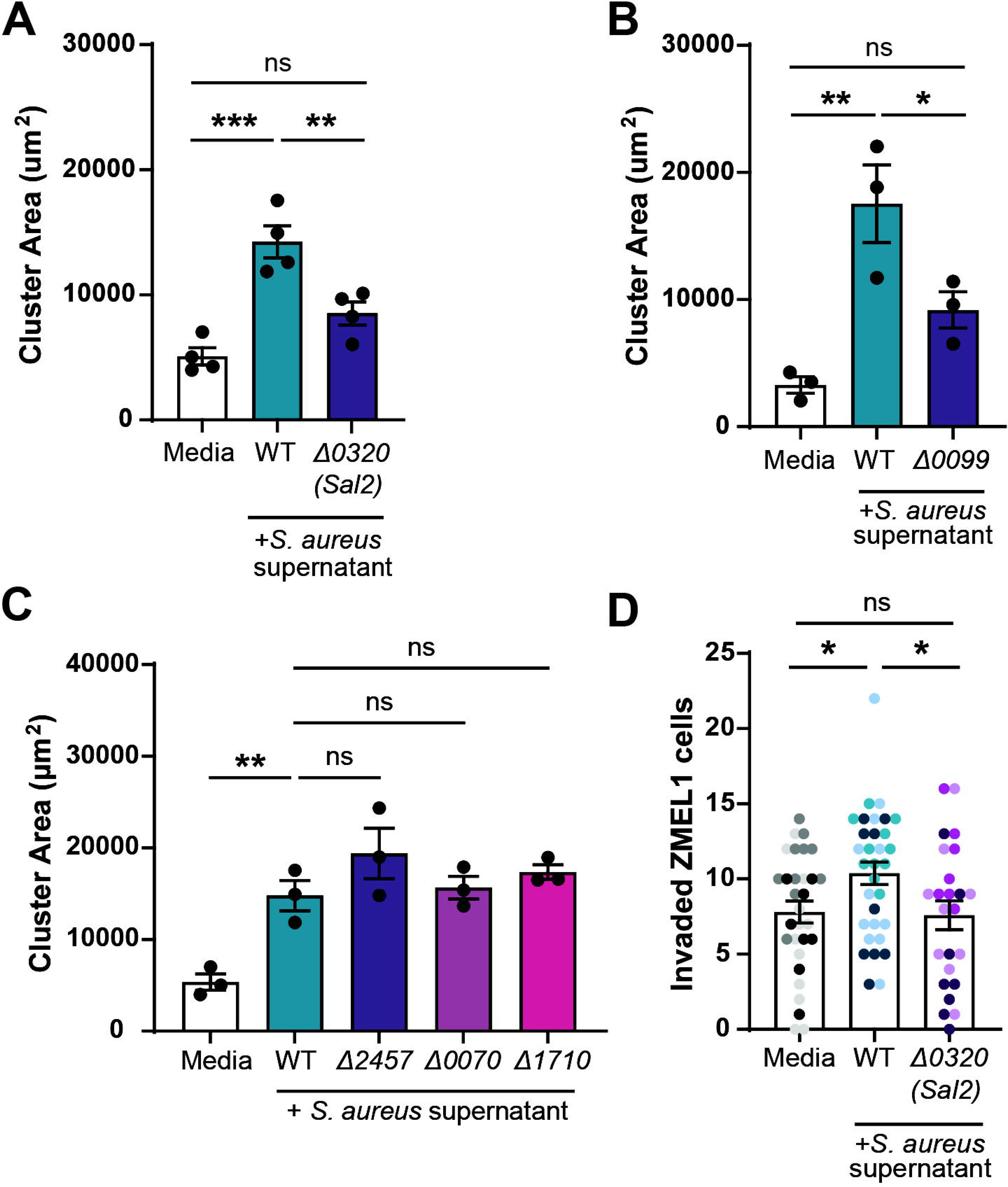
Deletion of *S. aureus* lipases alters melanoma clustering and invasion. Quantification of ZMEL1 melanoma cell cluster size (µm^2^) after 7 days of culture in media alone or plus *S. aureus* USA300 bacterial supernatant. Supernatants were tested from USA300 transposon mutants for (A) lipases *Sal2*/*Geh* (Δ0320) or (B) Phosphatidylinositol (PI)-specific phospholipase C (PI-PLC) (Δ*0099).* (C) Supernatants were tested from USA300 transposon mutants for putative phospholipase genes Δ2457, Δ0070, and Δ1710. (D) ZMEL1 invasion at 2dpi in the zebrafish hindbrain after culture in media alone or *S. aureus* USA300 supernatant. Dots represent independent larvae color coded per replicate. Media control n=30 larve, WT bacterial supernatant n=32 larvae, Δ0320 supernatant n=24 larvae. Experiments were conducted at least three times. Dots in (A-C) represent independent replicates and bars indicate the mean ± SEM. *p* values were calculated by One-way ANOVA (A-D). *, p < 0.05; **, p < 0.01; ***, p < 0.001.

To determine if the clustering effect was specific to SAL2, we also tested a selection of phospholipase mutants. Phosphatidylinositol (PI)-specific phospholipase C (PI-PLC) (SAUSA300_0099) is produced and secreted by all *S. aureus* strains, but most highly expressed by USA300 and Newman [46, 47]. Supernatant from the PI-PLC mutant Δ0099 revealed a significant decrease in clustering compared to WT (Figure 5B). Other putative phospholipase mutants (SAUSA300_2457, SAUSA300_0070, SAUSA300_1710) showed no significant effect on melanoma cluster size (Figure 5C). These findings suggest that bacterial lipid products mediate melanoma clustering.

Finally, we wanted to test the effect of these lipase mutants on melanoma behavior *in vivo*. We pre-incubated ZMEL1 cells with ΔSAL2 (Δ0320) *S. aureus* supernatant, injected the cells into the larval zebrafish hindbrain and imaged melanoma invasion at 2dpi. We found that melanoma cells incubated with this bacterial supernatant had a significant decrease in invasion compared to the WT supernatant (Figure 5D). Thus, *S. aureus* lipases likely produce lipids that induce melanoma cell clustering *in vitro* and invasion *in vivo*.

## Discussion

Changes in gut microbiota can affect cancer development, disease progression, and response to therapy [48]. The impact of the skin microbiota on carcinogenesis and cancer progression is less clear, but increasing evidence indicates that these microorganisms are also capable of influencing the balance between skin health and disease [49]. In this study, we found that *S. aureus* supernatant promotes melanoma invasion and dissemination in a transplant larval zebrafish model. Furthermore, *in vitro* analysis determined that lipids produced by *S. aureus* may be responsible for stimulating melanoma migration through Rho-GTPase activation.

Bacteria can promote cancer progression by impacting immune cell function in the tumor microenvironment [6]. Specifically, increased immune infiltration results in chronic or high-grade inflammation and induces tumor cell invasion and metastasis [5, 50]. We found that Zebrafish melanoma cells cultured with *S. aureus* bacterial supernatant were more migratory and invaded into the tissue *in vivo*. We quantified innate immune cell recruitment to determine if there was an increased inflammatory response to cancer cells that had been pre-treated with *S. aureus* bacterial supernatant, but we found no difference in neutrophil or macrophage numbers. Alternatively, *Fusobacterium nucleatum* promotes colorectal cancer cell migration by increasing secretion of inflammatory chemokines directly from the cancer cells themselves [6]. Melanoma cells express TLRs and initiate NFkB signaling, including production of chemotactic factors [36, 51, 52]. Thus, *S. aureus* may induce autocrine signaling of inflammatory chemokines to promote tumor cell migration.

Cancer metastasis is often associated with epithelial-to-mesenchymal transition (EMT) in which tumor cells acquire stem cell-like characteristics that enable invasion into the tissue. Microbes have been shown to directly activate signaling pathways involved in EMT [53-58]. In our study, melanoma cells pre-incubated with bacterial supernatant showed increased invasion and displayed a more mesenchymal morphology with increased dissemination to the tail fin. This phenotype was present despite washing the bacterial supernatant from melanoma cells before injection. I*n vitro* analysis determined that *S. aureus* secreted factors directly affect melanoma cell behavior. Melanoma clustering was regulated by Rho GTPase signaling and may similarly drive invasive cell migration *in vivo*.

Our work suggests that lipids produced by *S. aureus* are responsible for changes in melanoma cell behavior. Stripping of hydrophobic molecules with charcoal-dextran or Amberlite-XAD4 beads completely abolished any clustering effect of *S. aureus* supernatant. Furthermore, addition of bovine serum albumin increased cluster size 3-fold. Albumin proteins are known to bind serum lipids to increase cellular interactions with hydrophobic molecules in the blood [39, 40]. Taken together, it is likely that lipids secreted by *S. aureus* promote melanoma clustering and invasion. Only *S. aureus* strains USA300 and Newman were capable of promoting melanoma clustering, suggesting that these strains express specific lipid processing pathways not found in other gram-positive or-negative species tested. Thus, we evaluated the role of specific *S. aureus* lipase mutants on melanoma clustering.

*S. aureus* express a selection of extracellular secreted lipases including SAL2, also known as GehB (SAUSA300_0320). SAL2 has previously been shown to alter immune cell behavior by inactivating pathogen derived ligands [59]. For example, SAL2 cleaves esterified fatty acids on bacterial lipoproteins to prevent TLR2-mediated immune recognition and subsequent cytokine production [60]. In this study, mutation of *gehB* resulted in decreased melanoma clustering, suggesting SAL2 lipase activity produces lipid mediators that are recognized by melanoma. Phosphatidylinositol (PI)-specific phospholipase C (PI-PLC) may work in a similar manner, as mutation of *plc* resulted in decreased melanoma clustering. PI-PLC hydrolyzes membrane lipids such as PI and membrane protein anchors containing glycosyl-PI [47]. Furthermore, PI-PLC is highly expressed by *S. aureus* USA300 and Newman, but is not expressed by *S. epidermidis* [47], correlating to our clustering data. Cleavage of these membrane lipids may produce mediators capable of interacting with and modulating melanoma cells. It is widely accepted that bacterial lipids can be recognized by adaptive T and NK cells during antigen binding on the CD1 receptor [61]. Additionally, studies indicate they can be recognized by other eukaryotic and mammalian receptors including GPCRs [41, 62]. In the gut, short chain fatty acids (SCFAs) generated by bacterial fermentation can bind GPCRs to induce chemokine and cytokine production [63, 64].

Future studies will be needed to send these findings to human cancer cells. In addition, we did not evaluate the impact of live *S. aureus* bacteria. In previous work, *S. aureus* peptides were found bound to HLA class I and II molecules within human melanoma cells [65], indicating that these bacteria may gain entrance and influence melanoma transcriptional activity intracellularly, similar to *F. nucleatum* [6].

Many questions still remain about whether bacterial skin colonization leads to disease. For example, *S. aureus* was shown to activate mast cells and this heightened inflammation promotes the development of atopic dermatitis [66]. However, more data supports the theory that disruptions in the skin barrier lead to microbial dysbiosis [11, 67, 68]. Skin barrier disruption results in chronic inflammation and has been correlated with development of non-melanoma skin cancers [69, 70]. In cutaneous melanoma, ulceration is a negative prognostic factor and results in a damaged epidermal layer [71], which may allow for colonization of pathogenic species of *Staphylococcus*.

Here, we report that the skin microbe *S. aureus* promotes melanoma invasion and dissemination in a larval zebrafish tumor transplantation model. Our findings suggest that *S. aureus* lipase activity produces lipid mediators that modulate the behavior of melanoma cells. Our work supports further study on the role of bacterial-derived lipids on cancer cell invasion.

## Materials and Methods

### Ethics Statement

Animal care and use protocol M005405-A02 was approved by the University of Wisconsin-Madison College of Agricultural and Life Sciences (CALS) Animal Care and Use Committee. This protocol adheres to the federal Health Research Extension Act and the Public Health Service Policy on the Humane Care and Use of Laboratory Animals, overseen by the National Institutes of Health (NIH) Office of Laboratory Animal Welfare (OLAW).

### Fish Lines and Maintenance

Adult zebrafish and larvae (*Danio rerio)* were maintained as previously described [72]. Adult fish aged 3 months to 2 years were used to spawn larvae. Prior to experimental procedures, larvae were anesthetized in E3 water containing 0.2 mg/ml Tricaine (ethyl 3-aminobenzoate, Sigma). Larvae were maintained in E3 containing 0.2 mM N-phenylthiourea beginning 1 day post fertilization (dpf) (PTU, Sigma Aldrich) to prevent pigment formation during imaging experiments. All zebrafish lines used in this study are listed in **Supplemental Table 1.**

### Bacterial strains and growth conditions

Bacterial strains used in this study are described in **Supplemental Table 2**. *S. aureus* bacterial colonies were initiated on solid agar plates made with TSA (Sigma) or LB (Genesee) for other bacterial strains. Single colonies were picked and suspended in TSB or LB to initiate a liquid culture and were grown overnight with shaking at 37°C.

To generate bacterial supernatants, overnight cultures were diluted 1:100 into blank RPMI media, and grown with shaking at 225rpm until the optical density at 600nm (OD600) was approximately 0.9. For data in Figure 4A, we grew *S. aureus* in RPMI + 2%FBS or RPMI + 2%BSA. Cultures were centrifuged for 5 minutes at 3000xg and then filtered through a 0.2um SFCA filter to remove bacteria from the supernatant. Plated CFUs were used to normalize the filtered supernatant with the original culture media to 3e8 CFU/mL. Bacterial supernatant was aliquoted and frozen at −80°C for up to 3 months.

Mutant strains from the annotated Nebraska Transposon Mutant Library (NTML) generated in USA300 strain JE2 (BEI resources repository) were used in this study, **Supplemental Table 2.** For these strains, 2ug/mL erythromycin (Sigma) was used for antibiotic selection during overnight culture. Bacterial supernatants were generated in blank RPMI media. Prepared bacterial supernatants were utilized in zebrafish hindbrain injections or cluster formation assay as described below. The control USA300 JE2 strain was utilized for those experiments with transposon mutants (Figure 5A-D).

#### Charcoal/Amberlite stripping

*S. aureus* supernatant was stripped of lipids and polar molecules using Charcoal-dextran or Amberlite XAD4. Charcoal-dextran (Sigma C6241) was prepared as previously described [75]. Briefly, charcoal was added at a final concentration of 0.25% (w/v) to distilled water (pH 7.4) with added 0.25M sucrose, 1.5mM MgCl2, and 10mM HEPES and rotated overnight at 4°C. A volume of prepared charcoal-dextran was centrifuged at 500xg for 10 min to pellet the charcoal and the supernatant was aspirated. An equal volume of bacterial supernatant or blank RPMI media was added, vortexed to mix, and incubated for 12 hours at 4°C. The conical tube was centrifuged at 500xg for 10 min to pellet the charcoal.

Amberlite XAD4 (Thermo Fisher L14142.36) was alternatively utilized to remove polar molecules from bacterial supernatant, as previously described [76]. Amberlite beads (1% w/v) were measured out and washed with rotation in PBS for 3 hours at room temperature. The beads were allowed to settle to the bottom and the PBS media was removed. An equal volume of bacterial supernatant or blank RPMI media was added and incubated overnight with rotation at 4°C. The prepared supernatant was pipetted into a new tube, filtered with a 0.2um filter and frozen at −80°C for up to 3 months. Charcoal/Amberlite stripped supernatant was added to ZMEL1 cells for cluster assay as described below.

#### Heat Inactivation

*S. aureus* supernatant prepared as described above or *S. aureus* live bacteria were heated at 100°C for 15 minutes to denature/heat inactivate. Heat-inactivated bacteria were resuspended in blank RPMI media at 3e8 CFU/mL. Heat inactivated supernatant or heat-inactivated bacteria were added to ZMEL1 cells for cluster assay as described below.

### ZMEL1 cell culture

We utilized ZMEL1-EGFP or ZMEL1-tdTomato cell lines generated from a primary zebrafish melanoma model expressing BRAF V600E in a p53-/-background [82]. Cell lines were tested for mycoplasma contamination. ZMEL1 cells were cultured in 10% FBS-DMEM supplemented with 1% Glutamax and 1% penicillin-streptomycin on 10ug/mL fibronectin coated plates in a sterile 28°C incubator. To harvest, cells were washed with sterile PBS, trypsinized and then counted with a hemocytometer.

### Zebrafish ZMEL1 injection

Bacterial supernatants prepared in blank RPMI media were diluted 1:4 into ZMEL1 culture media and incubated on the ZMEL1 cells for 3 days. Cells were harvested, washed in PBS, then resuspended in HBSS media at a concentration of 8 x 10^7^ cells/mL. Cells were then loaded into thin-walled glass capillary injection needles. The needle was then calibrated to inject 1nl volume (15-20cells). Transgenic larvae were pre-screened for fluorescence using a zoomscope (EMS3/SyCoP3; Zeiss; Plan-NeoFluor Z objective). Anesthetized larvae were then placed on 3% Agarose plate made with E3 and microinjected with ZMEL1 cells, with time range set to “millisecond” and pressure set to ∼15PSI on the microinjector.

### Zebrafish Imaging and Quantification

#### Invasion assay

To assess ZMEL1 invasion at the injection site, 2 dpi (days post injection) larvae were anesthetized and mounted in a Z-wedgi [83] device such that the hindbrain was fully visible. Z-seried images (3.45 µm slices) of the hindbrain were acquired on a spinning disk confocal microscope (CSU-X; Yokogawa) with a confocal scan head on a Zeiss Observer Z.1 inverted microscope, Plan-Apochromat NA 0.8/20x objective, and a Photometrics Evolve EMCCD camera. Between imaging session larvae were kept in E3 with PTU in individual 24-well plates. Larvae that had tumor cells already separated from the injected cluster at 1hpi were excluded from the experiment.

#### Dissemination assay

At 4dpi, larvae were scored as the percentage of larvae that exhibited tumor cell dissemination to regions posterior to the first somite, outside of the spinal cord, as previously described [27]. Larvae were screened for tumor cells in the trunk or tail at 3 hours post injection to eliminate fish where ZMEL1 cells had been accidentally injected directly into the circulation. Screening and scoring of zebrafish larvae were done using a zoomscope (EMS3/SyCoP3; Zeiss; Plan-NeoFluor Z objective).

#### Neutrophil and Macrophage recruitment

Transgenic larvae were pre-screened for fluorescence using a zoomscope (EMS3/SyCoP3; Zeiss; Plan-NeoFluor Z objective). Larvae were anesthetized and mounted in a Z-wedgi [83] device such that the hindbrain was fully visible. Z-seried images (3.45 µm slices) of the hindbrain were acquired on a spinning disk confocal microscope (CSU-X; Yokogawa) with a confocal scan head on a Zeiss Observer Z.1 inverted microscope, Plan-Apochromat NA 0.8/20x objective, and a Photometrics Evolve EMCCD camera. For neutrophil recruitment, images were taken at 1 hour post injection (hpi) and 1 dpi. For macrophage recruitment, images were taken at 1 and 2 dpi.

#### Image analysis and processing

Images of larvae represent a 3-D rendering of the images generated on Imaris (v10.0). Invaded tumor cells were quantified as cells that were fully separated from the injected tumor cell cluster within the field of view. For neutrophil and macrophage recruitment, cells within 50 µm of the injected cluster were quantified.

### Cluster Formation Assay

Clusters were generated as previously described [25]. Briefly, ZMEL1 td-Tomato cells were harvested and resuspended in 10% FBS-DMEM supplemented with 1% Glutamax and 1% penicillin-streptomycin. RPMI blank media (40uL) or bacterial supernatant (40uL) was first added to an ultra-low attachment 96-well plate (Corning 3474). Then, 50,000 ZMEL1 cells/well were seeded in 120uL on top to mix. Plates were incubated at 28°C for up to 7 days to allow clusters to form. Plates were imaged on indicated days using an inverted fluorescent microscope (Nikon Eclipse TE300) with a 20× objective and an automated stage (Ludl Electronic Products) with a Prime BSI Express camera (Teledyne Photometrics). Environmental controls were set to 28°C with 5% CO_2_. Fluorescent images were analyzed using ImageJ software. All data were quantified from culture day 7, unless otherwise indicated.

For movies of cluster formation, starting on Day 5, ZMEL1 clusters were imaged every 30 minutes using the same imaging parameters described above. Videos were compiled using ImageJ software.

#### ROCK inhibitor

To block cell migration during cluster formation, we used ROCK inhibitor Y-27632 (BioTechne 1254). RPMI blank media or *S. aureus* USA300 supernatant was supplemented with inhibitor to yield a final concentration of 500nM. As described above, ZMEL1 cells were added on top to mix and incubated at 28°C for up to 7 days to allow clusters to form.

#### Pam3CSK4 and LPS

TLR agonists were added to melanoma cells for the cluster formation assay. Pam3CSK4 (Invivogen tlrl-pms) at final concentration 100ug/mL or LPS from *E. coli* (Sigma L2755) at final concentration 1ug/mL were added during setup of the cluster formation assay. As described above, ZMEL1 cells were added on top to mix and incubated at 28°C for up to 7 days to allow clusters to form.

#### Cluster quantification

The average cluster size of each image was quantified using ImageJ. Fluorescent EGFP images were processed as followed: *despeckled, auto thresholded,* converted to *binary*, and the *area of particles* greater than 30um were quantified. Any clusters on the edge of the image were excluded. A total of nine images were taken with three images per well and three wells per condition. At least 50 clusters were quantified per condition.

### Thin-layer chromatography

Lipids were extracted from 5mL of *S. aureus* USA300 bacterial supernatant samples via the Bligh and Dyer method [84] and dried under a stream of N_2_ gas. Lipids were resuspended in chloroform: methanol (2:1 v/v) and loaded onto a silica TLC plate (Sigma 60778). For nonpolar lipid separation, the plates were developed three times as in [85]: (1) hexane:MTBE:acetic acid (80:20:1) up to 50% of plate height; (2) hexane:benzene (1:1) up to 80% of plate height, then (3) hexane up to 90% of the plate. Plate was allowed to dry between each solvent system. For polar lipid separation, plates were developed using chloroform:methanol:water (65:25:4). Plates were stained in a sealed container of iodine gas for roughly 5 minutes. Lipid species were identified using the non-polar lipid mixture A (Cayman Chemical 29376) or polar lipid mixture (Cayman Chemical 29374).

### Statistics

All experiments with statistical analyses represent at least three independent replicates (N). Statistical significance was set to 0.05 and all statistical tests are two-tailed. The replicate number of zebrafish (n) for experiments in Figure 1C, D and Figure 5D is indicated in the figure legend. Larvae from a single clutch were randomly assigned to the control or experimental group. Zebrafish were excluded from analysis if no ZMEL1 cells could be identified in the hindbrain following injection. For zebrafish experiments, equivariance was checked using an F test and determined that the samples did meet the requirement. We tested for outliers using the robust regression and outlier removal (ROUT) method [86], but no outliers were identified. Analysis of ZMEL1 cell invasion in zebrafish larvae was performed on data pooled from 3 independent experiments. *p* values were calculated by unpaired T-test (Figure 1C) or One-way ANOVA with Tukey’s multiple comparisons (Figure 5D). Analysis of ZMEL1 dissemination in zebrafish larvae (Figure 1D) was performed on the percent of fish with dissemination using paired T-test.

Melanoma clustering with *S. aureus* supernatant versus heat inactivated bacteria, gram positive vs gram negative bacteria, and the NE USA300 transposon mutants was analyzed by One-way ANOVA with Tukey’s multiple comparisons. Melanoma clustering with *S. aureus* supernatant grown in the presence of FBS or BSA, or treatment with ROCK inhibitor, charcoal-dextran, or amberlite-XAD4 was analyzed by Two-way ANOVA with Tukey’s multiple comparisons (GraphPad Prism version 10). All graphical representations of data were created in GraphPad Prism version 10 and figures were ultimately assembled using Adobe Illustrator (Adobe version 23.0.6).

## Acknowledgements

We thank Dr. Lindsay Kalan for sharing bacterial skin isolates (LK869, LK894, LK494). We thank the lab of Dr. Richard White for sharing the ZMEL1 cell lines. We thank members of the Huttenlocher lab for helpful discussions about the manuscript.

## Competing Interests

None to disclose.

## Funding

Research reported in this publication was supported by the National Cancer Institute (NCI) of the National Institutes of Health (NIH) under Award Number R01-CA085862 and National Institute of Allergy and Infectious Diseases (NIAID) under Award Number T32AI055397. The content is solely the responsibility of the authors and does not necessarily represent the official views of the NIH. The funders had no role in study design, data collection and analysis, decision to publish, or preparation of the manuscript. The content is solely the responsibility of the authors and does not necessarily represent the official views of the National Institutes of Health.

## Data Availability

All relevant data are included in the manuscript.

## Author Contributions

M.A.G., G.R., L.S., and J.X.D. performed experiments. M.A.G., G.R., L.S., J.X.D., J.F., J.D.S., and A.H. analyzed and interpreted data. M.A.G. and A.H. designed the research and wrote the paper.

---

**Supplemental Table 1.**
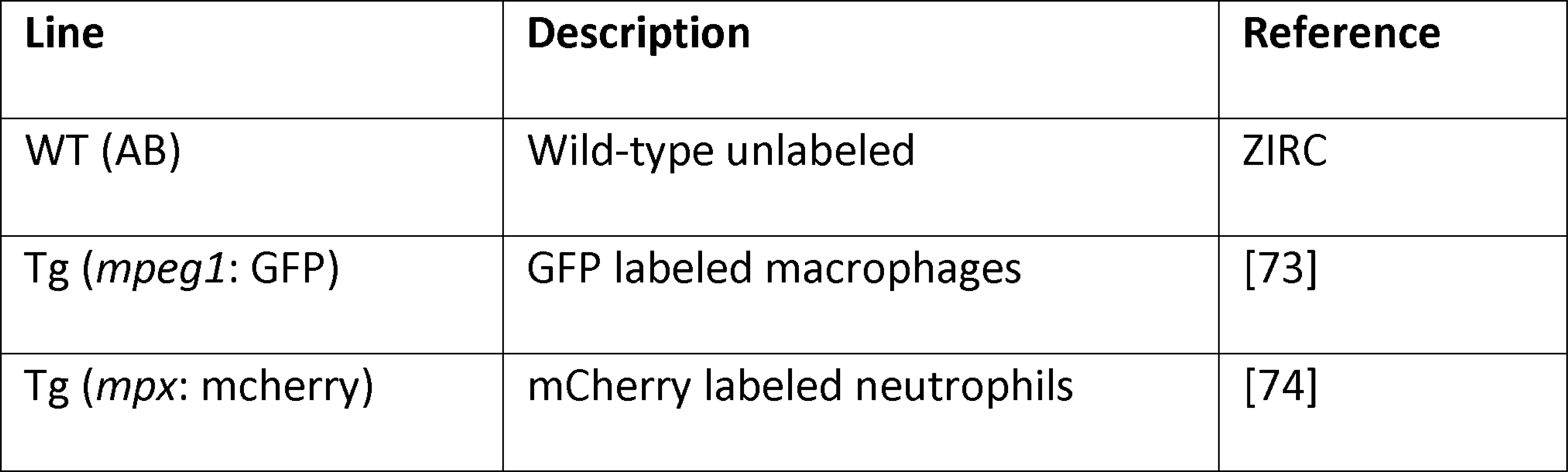
Zebrafish lines used in this study.

**Supplemental Table 2.**
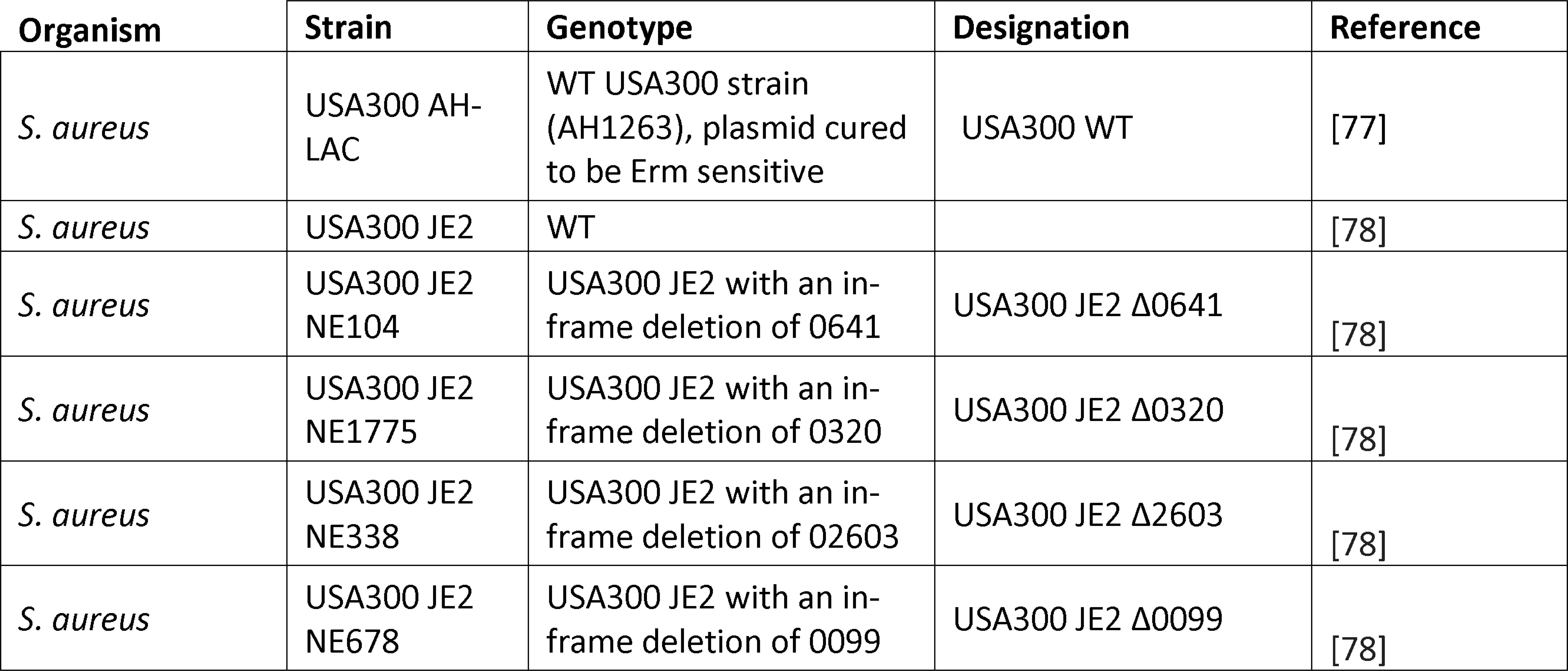
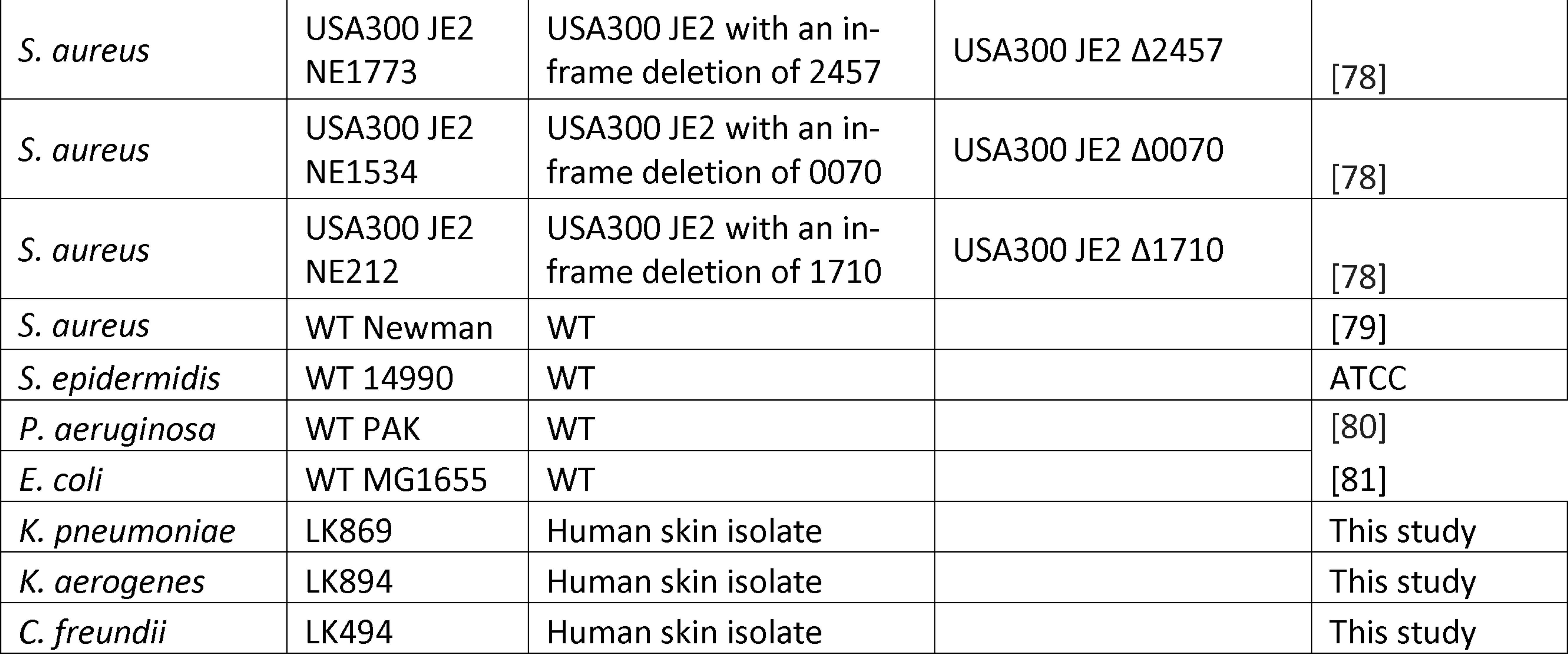
List of bacterial strains used in this study.

## Notes

### Competing Interest Statement

The authors have declared no competing interest.

## References

1. Zheng, D., T. Liwinski, and E. Elinav, Interaction between microbiota and immunity in health and disease. Cell Res, 2020. 30(6): p. 492–506.

2. Sun, J., F. Chen, and G. Wu, Potential effects of gut microbiota on host cancers: focus on immunity, DNA damage, cellular pathways, and anticancer therapy. ISME J, 2023. 17(10): p. 1535–1551.

3. Humans, I.W.G.o.t.E.o.C.R.t., Biological agents. IARC Monogr Eval Carcinog Risks Hum, 2012. 100(Pt B): p. 1–441.

4. Sepich-Poore, G.D., et al., The microbiome and human cancer. Science, 2021. 371(6536).

5. Garrett, W.S., Cancer and the microbiota. Science, 2015. 348(6230): p. 80–6.

6. Casasanta, M.A., et al., Fusobacterium nucleatum host-cell binding and invasion induces IL-8 and CXCL1 secretion that drives colorectal cancer cell migration. Sci Signal, 2020. 13(641).

7. Parhi, L., et al., Breast cancer colonization by Fusobacterium nucleatum accelerates tumor growth and metastatic progression. Nat Commun, 2020. 11(1): p. 3259.

8. Crusz, S.M. and F.R. Balkwill, Inflammation and cancer: advances and new agents. Nat Rev Clin Oncol, 2015. 12(10): p. 584–96.

9. Iida, N., et al., Commensal bacteria control cancer response to therapy by modulating the tumor microenvironment. Science, 2013. 342(6161): p. 967–70.

10. Gopalakrishnan, V., et al., Gut microbiome modulates response to anti-PD-1 immunotherapy in melanoma patients.Science, 2018. 359(6371): p. 97–103.

11. Yu, Y., et al., The role of the cutaneous microbiome in skin cancer: lessons learned from the gut. J Drugs Dermatol, 2015. 14(5): p. 461–5.

12. Squarzanti, D.F., et al., Non-Melanoma Skin Cancer: news from microbiota research. Crit Rev Microbiol, 2020. 46(4): p. 433–449.

13. Woo, Y.R., et al., The Human Microbiota and Skin Cancer. Int J Mol Sci, 2022. 23(3).

14. Swaney, M.H. and L.R. Kalan, Living in Your Skin: Microbes, Molecules, and Mechanisms. Infect Immun, 2021. 89(4).

15. Seidelman, J.L., C.R. Mantyh, and D.J. Anderson, Surgical Site Infection Prevention: A Review. JAMA, 2023. 329(3): p. 244–252.

16. Kullander, J., O. Forslund, and J. Dillner, Staphylococcus aureus and squamous cell carcinoma of the skin. Cancer Epidemiol Biomarkers Prev, 2009. 18(2): p. 472–8.

17. Wood, D.L.A., et al., A Natural History of Actinic Keratosis and Cutaneous Squamous Cell Carcinoma Microbiomes. mBio, 2018. 9(5).

18. Madhusudhan, N., et al., Molecular Profiling of Keratinocyte Skin Tumors Links Staphylococcus aureus Overabundance and Increased Human beta-Defensin-2 Expression to Growth Promotion of Squamous Cell Carcinoma. Cancers (Basel), 2020. 12(3).

19. Salava, A., et al., Skin microbiome in melanomas and melanocytic nevi. Eur J Dermatol, 2016. 26(1): p. 49–55.

20. Mekadim, C., et al., Dysbiosis of skin microbiome and gut microbiome in melanoma progression. BMC Microbiol, 2022. 22(1): p. 63.

21. Weiss, J.M., et al., Shifting the focus of zebrafish toward a model of the tumor microenvironment. Elife, 2022. 11.

22. Patton, E.E., et al., Melanoma models for the next generation of therapies. Cancer Cell, 2021. 39(5): p. 610–631.

23. White, R.M., et al., Transparent adult zebrafish as a tool for in vivo transplantation analysis. Cell Stem Cell, 2008. 2(2): p. 183–9.

24. Astell, K.R. and D. Sieger, Zebrafish In Vivo Models of Cancer and Metastasis. Cold Spring Harb Perspect Med, 2020. 10(8).

25. Campbell, N.R., et al., Cooperation between melanoma cell states promotes metastasis through heterotypic cluster formation. Dev Cell, 2021. 56(20): p. 2808–2825 e10.

26. Schoen, T.J., et al., Aspergillus fumigatus transcription factor ZfpA regulates hyphal development and alters susceptibility to antifungals and neutrophil killing during infection. PLoS Pathog, 2023. 19(5): p. e1011152.

27. Roh-Johnson, M., et al., Macrophage-Dependent Cytoplasmic Transfer during Melanoma Invasion In Vivo. Dev Cell, 2017. 43(5): p. 549–562 e6.

28. Barros-Becker, F., et al., Live imaging reveals distinct modes of neutrophil and macrophage migration within interstitial tissues. J Cell Sci, 2017. 130(22): p. 3801–3808.

29. Ribatti, D., R. Tamma, and T. Annese, Epithelial-Mesenchymal Transition in Cancer: A Historical Overview. Transl Oncol, 2020. 13(6): p. 100773.

30. Gonzalez, H., C. Hagerling, and Z. Werb, Roles of the immune system in cancer: from tumor initiation to metastatic progression. Genes Dev, 2018. 32(19-20): p. 1267–1284.

31. Giese, M.A., Hind, L.E., Huttenlocher, A., Neutrophil plasticity in the tumor microenvironment. Blood Rev, 2019.

32. Korte, B.G., et al., Cell Type-Specific Transcriptome Profiling Reveals a Role for Thioredoxin During Tumor Initiation. Front Immunol, 2022. 13: p. 818893.

33. Amano, M., M. Nakayama, and K. Kaibuchi, Rho-kinase/ROCK: A key regulator of the cytoskeleton and cell polarity. Cytoskeleton (Hoboken), 2010. 67(9): p. 545–54.

34. Jeong, K.J., et al., The Rho/ROCK pathway for lysophosphatidic acid-induced proteolytic enzyme expression and ovarian cancer cell invasion. Oncogene, 2012. 31(39): p. 4279–89.

35. Duan, T., et al., Toll-Like Receptor Signaling and Its Role in Cell-Mediated Immunity. Front Immunol, 2022. 13: p. 812774.

36. Burns, E.M. and N. Yusuf, Toll-like receptors and skin cancer. Front Immunol, 2014. 5: p. 135.

37. Baba, T., et al., Genome sequence of Staphylococcus aureus strain Newman and comparative analysis of staphylococcal genomes: polymorphism and evolution of two major pathogenicity islands. J Bacteriol, 2008. 190(1): p. 300–10.

38. Hines, K.M., et al., Lipidomic and Ultrastructural Characterization of the Cell Envelope of Staphylococcus aureus Grown in the Presence of Human Serum. mSphere, 2020. 5(3).

39. van der Vusse, G.J., Albumin as fatty acid transporter. Drug Metab Pharmacokinet, 2009. 24(4): p. 300–7.

40. Francis, G.L., Albumin and mammalian cell culture: implications for biotechnology applications. Cytotechnology, 2010. 62(1): p. 1–16.

41. Ryan, E., S.A. Joyce, and D.J. Clarke, Membrane lipids from gut microbiome-associated bacteria as structural and signalling molecules. Microbiology (Reading), 2023. 169(3).

42. Sohlenkamp, C. and O. Geiger, Bacterial membrane lipids: diversity in structures and pathways. FEMS Microbiol Rev, 2016. 40(1): p. 133–59.

43. Cadieux, B., et al., Role of lipase from community-associated methicillin-resistant Staphylococcus aureus strain USA300 in hydrolyzing triglycerides into growth-inhibitory free fatty acids. J Bacteriol, 2014. 196(23): p. 4044–56.

44. Altschul, S.F., et al., Basic local alignment search tool. J Mol Biol, 1990. 215(3): p. 403–10.

45. Rosenstein, R. and F. Gotz, Staphylococcal lipases: biochemical and molecular characterization. Biochimie, 2000. 82(11): p. 1005–14.

46. White, M.J., et al., Phosphatidylinositol-specific phospholipase C contributes to survival of Staphylococcus aureus USA300 in human blood and neutrophils. Infect Immun, 2014. 82(4): p. 1559–71.

47. Volwerk, J.J., et al., Phosphatidylinositol-specific phospholipase C from Bacillus cereus combines intrinsic phosphotransferase and cyclic phosphodiesterase activities: a 31P NMR study. Biochemistry, 1990. 29(35): p. 8056–62.

48. Hanahan, D., Hallmarks of Cancer: New Dimensions. Cancer Discov, 2022. 12(1): p. 31–46.

49. Savoia, P., et al., Role of the Microbiota in Skin Neoplasms: New Therapeutic Horizons. Microorganisms, 2023. 11(10).

50. Smith, C.K. and G. Trinchieri, The interplay between neutrophils and microbiota in cancer. J Leukoc Biol, 2018.

51. Sharma, B., et al., Targeting CXCR1/CXCR2 receptor antagonism in malignant melanoma. Expert Opin Ther Targets, 2010. 14(4): p. 435–42.

52. Wang, J.M., et al., Induction of haptotactic migration of melanoma cells by neutrophil activating protein/interleukin-8. Biochem Biophys Res Commun, 1990. 169(1): p. 165–70.

53. Arthur, J.C., et al., Intestinal inflammation targets cancer-inducing activity of the microbiota. Science, 2012. 338(6103): p. 120–3.

54. Liu, J., et al., Current Understanding of Microbiomes in Cancer Metastasis. Cancers (Basel), 2023. 15(6).

55. Vincan, E. and N. Barker, The upstream components of the Wnt signalling pathway in the dynamic EMT and MET associated with colorectal cancer progression. Clin Exp Metastasis, 2008. 25(6): p. 657–63.

56. Rubinstein, M.R., et al., Fusobacterium nucleatum promotes colorectal carcinogenesis by modulating E-cadherin/beta-catenin signaling via its FadA adhesin. Cell Host Microbe, 2013. 14(2): p. 195–206.

57. Lu, R., et al., Enteric bacterial protein AvrA promotes colonic tumorigenesis and activates colonic beta-catenin signaling pathway. Oncogenesis, 2014. 3(6): p. e105.

58. Franco, A.T., et al., Activation of beta-catenin by carcinogenic Helicobacter pylori. Proc Natl Acad Sci U S A, 2005. 102(30): p. 10646–51.

59. Stoll, H., et al., Staphylococcus aureus deficient in lipidation of prelipoproteins is attenuated in growth and immune activation. Infect Immun, 2005. 73(4): p. 2411–23.

60. Chen, X. and F. Alonzo, 3rd, Bacterial lipolysis of immune-activating ligands promotes evasion of innate defenses. Proc Natl Acad Sci U S A, 2019. 116(9): p. 3764–3773.

61. Gras, S., et al., Molecular recognition of microbial lipid-based antigens by T cells. Cell Mol Life Sci, 2018. 75(9): p. 1623–1639.

62. Cohen, L.J., et al., Commensal bacteria make GPCR ligands that mimic human signalling molecules. Nature, 2017. 549(7670): p. 48–53.

63. Tan, J., et al., The role of short-chain fatty acids in health and disease. Adv Immunol, 2014. 121: p. 91–119.

64. Chun, E., et al., Metabolite-Sensing Receptor Ffar2 Regulates Colonic Group 3 Innate Lymphoid Cells and Gut Immunity. Immunity, 2019. 51(5): p. 871–884 e6.

65. Kalaora, S., et al., Identification of bacteria-derived HLA-bound peptides in melanoma. Nature, 2021. 592(7852): p. 138–143.

66. Nakamura, Y., et al., Staphylococcus delta-toxin induces allergic skin disease by activating mast cells. Nature, 2013. 503(7476): p. 397–401.

67. Zeeuwen, P.L., et al., Microbiome dynamics of human epidermis following skin barrier disruption. Genome Biol, 2012. 13(11): p. R101.

68. Grice, E.A. and J.A. Segre, Interaction of the microbiome with the innate immune response in chronic wounds. Adv Exp Med Biol, 2012. 946: p. 55–68.

69. Williams, M.R., et al., Interplay of Staphylococcal and Host Proteases Promotes Skin Barrier Disruption in Netherton Syndrome. Cell Rep, 2020. 30(9): p. 2923–2933 e7.

70. Hoste, E., et al., Innate sensing of microbial products promotes wound-induced skin cancer. Nat Commun, 2015. 6: p. 5932.

71. Balch, C.M., et al., Final version of 2009 AJCC melanoma staging and classification. J Clin Oncol, 2009. 27(36): p. 6199–206.

72. Vincent, W.J., et al., Macrophages mediate flagellin induced inflammasome activation and host defense in zebrafish. Cell Microbiol, 2016. 18(4): p. 591–604.

73. Ellett, F., et al., mpeg1 promoter transgenes direct macrophage-lineage expression in zebrafish. Blood, 2011. 117(4): p. e49–56.

74. Yoo, S.K., et al., Differential regulation of protrusion and polarity by PI3K during neutrophil motility in live zebrafish. Dev Cell, 2010. 18(2): p. 226–36.

75. Sikora, M.J., et al., Endocrine Response Phenotypes Are Altered by Charcoal-Stripped Serum Variability. Endocrinology, 2016. 157(10): p. 3760–3766.

76. Hansen, S., et al., A Novel Growth-Based Selection Strategy Identifies New Constitutively Active Variants of the Major Virulence Regulator PrfA in Listeria monocytogenes. J Bacteriol, 2020. 202(11).

77. Miller, L.G., et al., Necrotizing fasciitis caused by community-associated methicillin-resistant Staphylococcus aureus in Los Angeles. N Engl J Med, 2005. 352(14): p. 1445–53.

78. Fey, P.D., et al., A genetic resource for rapid and comprehensive phenotype screening of nonessential Staphylococcus aureus genes. mBio, 2013. 4(1): p. e00537–12.

79. Duthie, E.S. and L.L. Lorenz, Staphylococcal coagulase; mode of action and antigenicity. J Gen Microbiol, 1952. 6(1-2): p. 95–107.

80. Takeya, K. and K. Amako, A rod-shaped Pseudomonas phage. Virology, 1966. 28(1): p. 163-5.

81. Panagiotidis, C.A., et al., Biosynthesis of polyamines in ornithine decarboxylase, arginine decarboxylase, and agmatine ureohydrolase deletion mutants of Escherichia coli strain K-12. Proc Natl Acad Sci U S A, 1987. 84(13): p. 4423–7.

82. Heilmann, S., et al., A Quantitative System for Studying Metastasis Using Transparent Zebrafish. Cancer Res, 2015. 75(20): p. 4272–4282.

83. Huemer, K., et al., zWEDGI: Wounding and Entrapment Device for Imaging Live Zebrafish Larvae. Zebrafish, 2017. 14(1): p. 42–50.

84. Bligh, E.G. and W.J. Dyer, A rapid method of total lipid extraction and purification. Can J Biochem Physiol, 1959. 37(8): p. 911–7.

85. Choa, R., et al., Thymic stromal lymphopoietin induces adipose loss through sebum hypersecretion. Science, 2021. 373(6554).

86. Motulsky, H.J. and R.E. Brown, Detecting outliers when fitting data with nonlinear regression - a new method based on robust nonlinear regression and the false discovery rate. BMC Bioinformatics, 2006. 7: p. 123.

